# Machine Learning Methods to Identify Genetic Correlates of Radiation-Associated Contralateral Breast Cancer in the WECARE Study

**DOI:** 10.1101/547422

**Authors:** Sangkyu Lee, Xiaolin Liang, Meghan Woods, Anne S. Reiner, Duncan Thomas, Patrick Concannon, Leslie Bernstein, Charles F. Lynch, John D. Boice, Joseph O. Deasy, Jonine L. Bernstein, Jung Hun Oh

**Affiliations:** Department of Medical Physics, Memorial Sloan Kettering Cancer Center, New York, NY; Department of Epidemiology and Biostatistics, Memorial Sloan Kettering Cancer Center, New York, NY; Department of Preventive Medicine, University of Southern California, Los Angeles, CA; Genetics Institute and Department of Pathology, Immunology and Laboratory Medicine, University of Florida, Gainesville, FL; Department of Population Sciences, Beckman Research Institute of the City of Hope, Duarte, CA; Department of Epidemiology, The University of Iowa, Iowa City, IA; Department of Medicine, Vanderbilt University Medical Center, Nashville, TN

## Abstract

The purpose of this study is to identify germline single nucleotide polymorphisms (SNPs) that optimally predict radiation-associated contralateral breast cancer (RCBC) and to provide new biological insights into the carcinogenic process. Fifty-two women with contralateral breast cancer and 153 women with unilateral breast cancer were identified within the Women’s Environmental Cancer and Radiation Epidemiology (WECARE) Study who were at increased risk of RCBC because they were ≤ 40 years of age at first diagnosis of breast cancer and received a scatter radiation dose > 1 Gy to the contralateral breast. A previously reported algorithm, preconditioned random forest regression, was applied to predict the risk of developing RCBC. The resulting model produced an area under the curve of 0.62 (*p*=0.04) on hold-out validation data. The biological analysis identified the cyclic AMP-mediated signaling and Ephrin-A as significant biological correlates, which were previously shown to influence cell survival after radiation in an ATM-dependent manner. The key connected genes and proteins that are identified in this analysis were previously identified as relevant to breast cancer, radiation response, or both. In summary, machine learning/bioinformatics methods applied to genome-wide genotyping data have great potential to reveal plausible biological correlates associated with the risk of RCBC.

## Introduction

Radiation-associated contralateral breast cancer (RCBC) is a rare adverse health outcome following radiation therapy for primary breast cancer; young women at the time of exposure are at greater risk than older women ^1–4^. A number of risk or modifying factors have been found to be associated with developing contralateral breast cancer (CBC) such as the number of full-term pregnancies ^5^, age at menarche ^6–8^, treatment with chemotherapy or duration of tamoxifen therapy for first breast cancer ^9–11^, family history of breast cancer ^12–14^, and radiotherapy dose ^3,15^. Genetic factors also have been identified ^16–20^ suggesting that certain rare genetic variations in deoxyribonucleic acid (DNA) damage response genes such as ATM might make it difficult to repair the damage induced by radiation and eventually may lead to CBC. Germline variations in genes associated with damage response also have been reported as breast cancer susceptibility loci among the general population ^21–23^. However, the candidate gene approach used in these studies has not provided new biological insights beyond the already known DNA damage response such as DNA repair or cell cycle arrest.

Genome-wide association studies (GWAS) — agnostic testing of associations between a phenotype and common single nucleotide polymorphisms (SNPs) sampled across the genome — can complement the candidate gene approach in discovering new genes associated with CBC. For example, GWAS have identified loci that confer risk for breast cancer ^24^ as well as radiation-associated thyroid cancer ^25^ and malignancies following radiotherapy for lymphoma ^26^. Recently, it was found that common breast cancer risk loci from GWAS ^24^ are related to CBC risk in a manner consistent with a polygenic risk score (PRS) ^27^. However, a gap in understanding is the contribution to CBC risk of biological factors other than the known breast cancer or DNA response genes and any interactions between SNPs that are not accounted for in a PRS. When applied to GWAS data, an agnostic non-linear modeling approach could fill this gap, thereby potentially enabling the discovery of new biomarkers associated with CBC and improving the ability to predict the risk of CBC following radiotherapy.

The specific goal of this study was to apply an agnostic machine learning approach to GWAS data to identify a predictive risk model for RCBC. A secondary goal was to gain novel biological insights from the predictive model using a systematic bioinformatics approach and a knowledge mining method from curated biological databases. In the current study, the focus is on a subgroup of women who received scatter and leakage radiation dose > 1 Gy to the contralateral breast at a young age (≤ 40 years) from the Women’s Environmental Cancer and Radiation Epidemiology (WECARE) Study ^28^. A machine-learning/bioinformatics methodology was employed, which was previously used to model radiation-induced complications of late rectal bleeding and erectile dysfunction ^29^, and chronic urinary dysfunction following radiotherapy ^30^.

## Materials and Methods

### Study population

The WECARE Study is a multi-center, population-based case-control study of breast cancer survivors designed to evaluate CBC risk following breast cancer treatments. Data from the first phase of the WECARE Study, WECARE I ^28^ was used; participants were recruited from five cancer registries in the United States and Denmark: Surveillance Epidemiology and End Results (SEER) registries in Iowa; Seattle, Washington; Los Angeles County, California; Orange County, California; and the Danish Breast Cancer Cooperative Group registry. Ethical approval was obtained at each local site. All patients provided informed consent and the study protocols were approved by the institutional review boards at each US site including University of Iowa, Fred Hutchinson Cancer Research Center, University of Southern California, and University of California at Irvine, and by the Ethics Committee System in Denmark. All experiments were performed in accordance with relevant guidelines.

The cases were women who had a first primary breast cancer diagnosis and developed CBC one or more years later. The controls were women with primary breast cancer who did not develop CBC after having been followed for at least as long as their matched case. The study participants were genotyped using the Illumina HumanOmni1-Quad version 1.0, reporting 822,778 germline SNPs. Quality control filters were used, including minor allele frequency (MAF) ≥ 0.01, missing rate ≤ 0.05 for SNPs and individuals, and Hardy-Weinberg Equilibrium (HWE) *p* > 10^-5^. The genotyped SNPs were imputed for sporadic missing genotypes using IMPUTE2 software and the 1000 Genomes reference panel ^31, 32^. As a result, 767,207 SNPs with the resulting imputation probability > 0.9 were used for further analysis.

Because analyses of data from the WECARE Study showed that CBC risk is greater in younger women who received contralateral breast doses > 1 Gy, the cohort of this study was restricted to women (cases and controls) who met the following criteria: 1) Caucasian with European origin, 2) age ≤ 40 years at first breast cancer diagnosis, and 3) radiation scatter dose to the contralateral breast > 1 Gy (dosimetric procedures are described in ^15^)). These criteria resulted in 205 eligible women (52 cases and 153 controls) selected from the WECARE I Study of 2,102 women (705 cases and 1,397 controls). The study sample was randomly split into training (N = 137) and validation (N = 68) sets to rigorously test the results. The training and validation datasets were balanced with respect to case-control ratio, age at diagnosis, average radiation dose to the contralateral breast, and cancer registry; in dividing the dataset, priority was given in the order listed because a perfect split could not be achieved for all of the variables (Table 1).

**Table 1.** Patient characteristics of training and validation datasets.

|  | Training | Validation |
| --- | --- | --- |
| Sample size | 137 | 68 |
| Control | 102 (74.4%) | 51 (75.0%) |
| Case | 35 (25.6%) | 17 (25.0%) |
| Age at diagnosis (years) |  |  |
| Median | 37 | 37 |
| Minimum | 23 | 24 |
| Maximum | 40 | 40 |
| Average radiation dose to the contralateral breast (Gy) | 1.32 | 1.37 |
| Median | 1.01 | 1.01 |
| Minimum | 3.74 | 2.90 |
| Maximum |  |  |
| Registry |  |  |
| Denmark | 16 (11.7%) | 7 (10.3%) |
| Iowa | 32 (23.4%) | 16 (23.5%) |
| Orange County, California | 24 (17.5%) | 11 (16.2%) |
| Los Angeles County, California | 57 (41.6%) | 28 (41.2%) |
| Seattle, Washington | 8 (5.8%) | 6 (8.8%) |

### Univariate analysis of SNPs and clinical variables

Univariate associations in the training set between RCBC status and all the candidate predictors, including SNPs and clinical variables, were examined. For the SNPs, the association was tested using the chi-square test under an additive model. Based on prior analyses within the WECARE I Study, we evaluated the following 6 clinical variables: family history of breast cancer ^33^, number of full-term pregnancies ^5^, age at menarche ^5, 6^, treatment with chemotherapy or duration of tamoxifen therapy for first breast cancer ^34^, age at diagnosis of the first breast cancer, and scatter radiation dose to the contralateral breast ^15^. The statistical significance of variability in RCBC rates among the 5 registry locations was tested (Table 2). To assess the difference between cases and controls, categorical and continuous variables were tested using the chi-square test and univariate logistic regression, respectively. To reduce high-dimensionality burden to predictive model training, a filtering approach was employed where SNPs were removed based on univariate association strength prior to the modeling ^35^: SNPs with association *p*-values < 0.001 and the clinical variables with association *p*-values < 0.05 were incorporated into predictive model building. The univariate *p*-value cutoff value was taken as in previous studies ^29, 30^.

**Table 2.** Statistical significance of associations between radiation-associated contralateral breast cancer and clinical variables in the training dataset of the study cohort. Numbers in parentheses: 95% confidence intervals for odds ratios.

| Variables | Levels | Odds ratio | *P*-value |
| --- | --- | --- | --- |
| Age at menarche (years) | < 13 (Reference)<br>≥ 13 | 0.98 (0.42, 2.27) | 1.00 |
| Family history of breast cancer | None (Reference)<br>1+ | 1.78 (0.70, 4.44) | 0.25 |
| Chemotherapy or hormonal treatment | None (Reference)<br>Yes | 0.80 (0.30, 2.22) | 0.79 |
| Number of full-term pregnancies | None (Reference)<br>1<br>2<br>3<br>>3 | 1.60 (0.48, 5.29)<br>1.01 (0.34, 3.03)<br>1.35 (0.25, 6.13)<br>1.68 (0.03, 35.50) | 0.88 |
| Age at first breast cancer diagnosis | Continuous | 1.02 (0.91, 1.13) | 0.76 |
| Average radiation dose to the contralateral breast | Continuous | 1.47 (0.69, 3.11) | 0.32 |
| Registry | Denmark (Reference)<br>Iowa<br>Orange County, California<br>Los Angeles County, California<br>Seattle, Washington | 0.42 (0.08, 2.20)<br>0.74 (0.15, 3.86)<br>0.94 (0.25, 3.98)<br>0.74 (0.05, 6.55) | 0.65 |

### RCBC prediction model training and validation

Using the reduced set of SNPs, we trained a multivariate model for predicting RCBC status based on genotypes and clinical characteristics. Specifically, the Preconditioned Random Forest Regression (PRFR) method was used as the learning strategy (see ^29^ and ^30^). In brief, an initial preconditioning task transforms the original binary outcomes (1: occurrence of RCBC, 0: no occurrence) into continuous ([0,1]) probability-based estimations of outcome risk (preconditioned outcomes) by regressing the original outcomes on supervised principal components of the most highly associated SNPs (note that this transformation only takes place for outcomes in the training set; the outcomes in the validation set are retained as binary for evaluation of classification accuracy).

Subsequently, a random forest (RF) model was built to optimally predict the preconditioned outcomes with respect to the predictors (SNPs). The RF is a multivariate method that consists of an ensemble (forest) of decision trees. Each tree is built with a bootstrap subsample of the training data. It is expected that these types of trees are well suited to consider non-linear biological dependencies. RF has been used for GWAS because of its flexibility in accounting for non-linear effects between SNPs and the robustness in high-dimensional problems ^36^. Each tree produces a predicted probability for a validation sample at a terminal leaf in the following way: each validation sample passes down through the built tree and reaches to a terminal leaf. The average preconditioned outcome of the samples in the terminal leaf is used as prediction for that tree. The overall prediction for the sample is then the average prediction across the whole forest.

To assess predictive power, the area under the curve (AUC) was calculated between the binary outcomes and the probability of RCBC, as predicted by the PRFR model. Performance of the PRFR was compared against other multivariate models such as conventional RF classification and least absolute shrinkage and selection operator (LASSO) models; variability of predictive performance on the validation dataset was assessed and compared between the modeling approaches. To this end, a 5-fold process using 80% of the training set was repeated 100 times, with random data shuffling between runs, resulting in 500 models. Each model was then applied to the hold-out validation data, resulting in the range of validation AUCs.

### SNP prioritization and identification of biological correlates

The impact of individual SNPs on model prediction of RCBC was assessed by a variable importance measure (VIM). The VIM was computed through the RF modeling for each predictor to quantify its contribution to prediction accuracy. To compute VIM, a permutation-based approach was used, where the values of each predictor were shuffled across samples while keeping other predictors fixed. The resulting increase in prediction mean squared error, due to loss of predictive information caused by the shuffling, was calculated using out-of-bag samples. We used the VIM metric as a secondary filtering approach to complement the preceding univariate test to further isolate the SNPs that are more important in predicting RCBC and thus more likely to be biologically relevant ^37^. Thus, we examined biological correlates using the SNPs in the top 25%, 50%, and 75% of VIM that maximized the validation AUC acquired using 80% of the training set.

For the biological analysis, SNPs were mapped to genes by including the genes that are located within 50,000 base pairs of each SNP. Gene and SNP coordinates were taken from the human genome build 19 (hg19). Using the resulting gene list, the enrichment of biological process terms in the Gene Ontology (GO) annotation database was investigated ^38^. The ClueGO software ^39^ was used to perform a gene set enrichment analysis (GSEA) and to organize the significant processes into relevant groups based on the number of common annotated genes. Details of the GSEA methodology can be found in the Supplementary material of our previous work ^30^. GO terms with a false discovery rate (FDR) *p* ≤ 0.05 were reported. We also searched for a cluster of interacting gene products from the list to identify those most likely to manifest similar biological functions. MetaCore^TM^ (Thompson Reuters, New York, NY) was used for mining previously known interactions between the queried gene products. A systematic literature survey was conducted on the gene products that formed the clusters. The search intended to test two independent hypotheses: 1) these markers are involved in breast cancer carcinogenesis independently of radiation, and 2) these markers are known to respond to radiation that increases breast cancer risk. To this end, the following keywords were searched for within PubMed for each hypothesis: 1) “breast neoplasms” [MeSH Terms] AND (*candidate protein name*), or 2) (“radiation, ionizing” [MeSH Terms] OR “ultraviolet rays” [MeSH Terms]) AND (“neoplasms” [MeSH Terms] OR “carcinogenesis” [MeSH Terms]) AND (*candidate protein name*).

Moreover, the impact of different ways in prioritizing SNPs on biological interpretation was investigated. In addition to the VIM-based ranking from the PRFR model, we considered: 1) the relative SNP importance defined as the frequency of selection by the LASSO model over 100 iterations, and 2) SNP ranking by univariate association *p-*values. The GSEA results were compared between the three SNP prioritization methods keeping the number of genes used for the GSEA fixed at the same number as in the VIM-based ranking.

All statistical analyses were conducted using the R language-based statistical libraries. A simplified description of the data analysis pipeline is shown in Figure 1.

**Figure 1:**
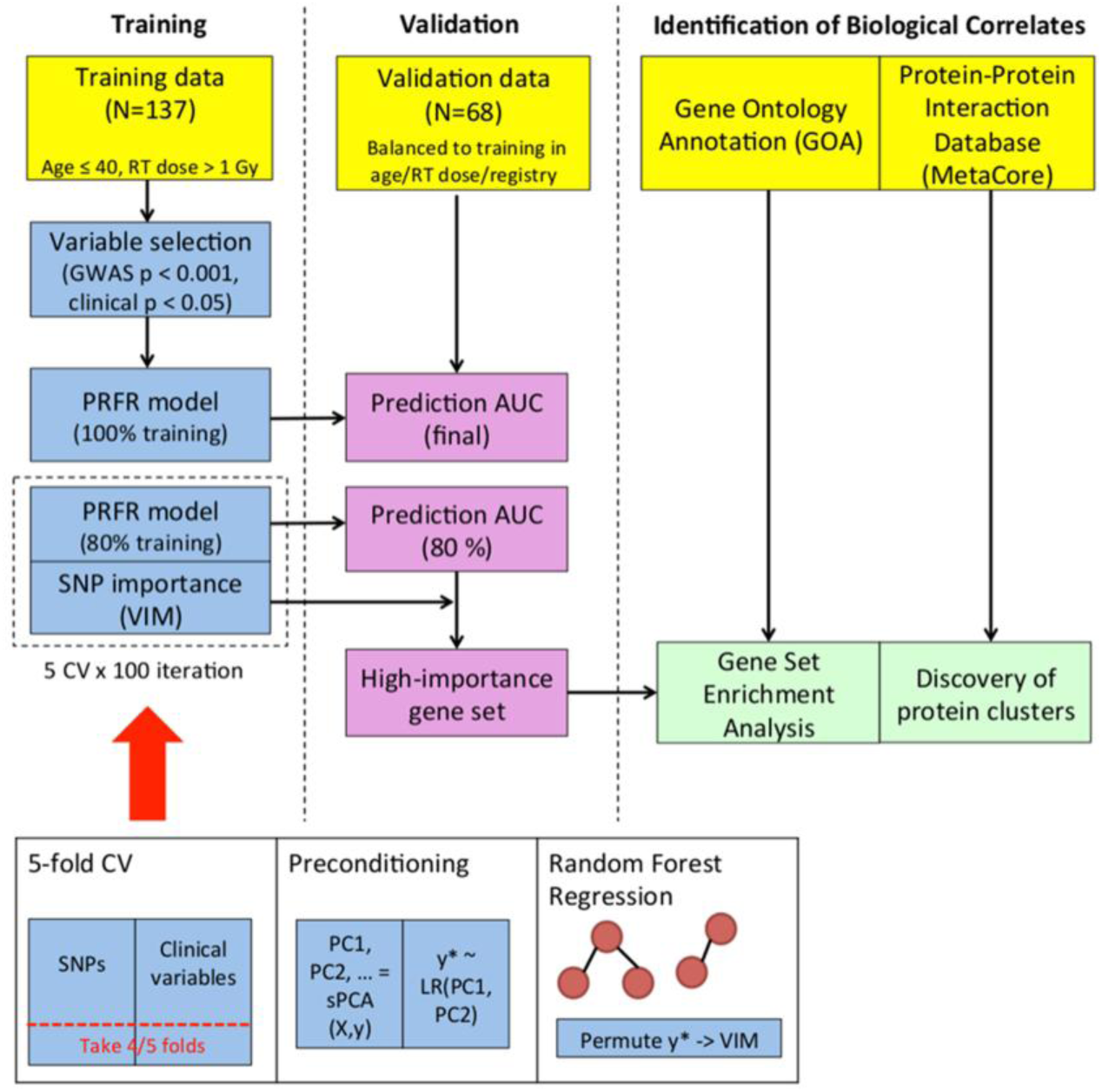
A flowchart summarizing the data analysis pipeline. *Abbreviations*: RT - radiotherapy; PC - principal component; sPCA - supervised principal component analysis; LR - logistic regression; GWAS - genome-wide association studies; AUC - area under the curve; CV - cross-validation; PRFR - preconditioned random forest regression; SNP - single nucleotide polymorphism; VIM - variable importance measure.

## Results

### Statistical significance of the genome-wide SNPs and clinical factors

No notable inflation (λ~0.994) from the *p-*values of the genome-wide associations was found (Figure 2), indicating negligible evidence of population stratification ^40^. The lowest univariate *p*-value was observed for rs2298515 (*p* = 5.6 × 10^-7^). In the initial filtering using a *p*-value cutoff of 0.001, 712 SNPs were left as predictors for the modeling steps. No clinical variables, including registry, reached nominal significance (*p* < 0.05) (Table 2). Thus, the subsequent PRFR model was built using only the 712 SNP predictors.

**Figure 2:**
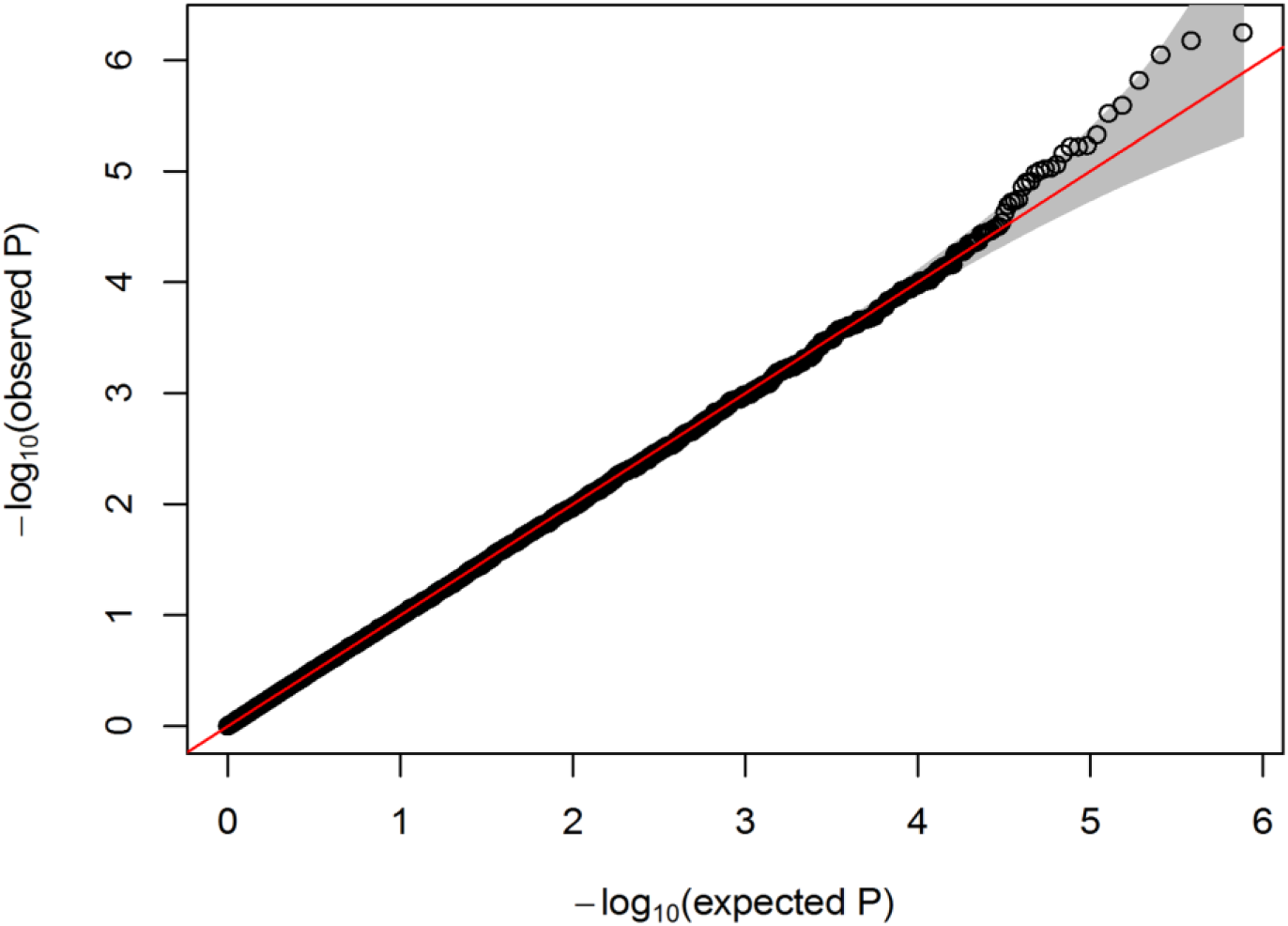
A quantile-quantile plot indicating association *p*-values between 767,207 single nucleotide polymorphisms and case/control status of contralateral breast cancer.

### Predictive performance of the PRFR model

When 80% of the training data was used, the PRFR model achieved an average validation AUC of 0.57 (95% confidence interval [CI]: 0.57 - 0.58). This was statistically significantly (t-test *p* < 0.001) higher than the AUCs for the conventional RF (AUC = 0.55, 95% CI: 0.55 - 0.56), LASSO (AUC = 0.55, 95% CI: 0.55 - 0.56), or preconditioned LASSO (AUC = 0.53, 95% CI: 0.53 - 0.54) (Figure 3). When the PRFR was retrained using all the available training set, the validation AUC increased to 0.62 (bootstrap estimated 95% CI: 0.48 - 0.75, *p =* 0.04).

**Figure 3:**
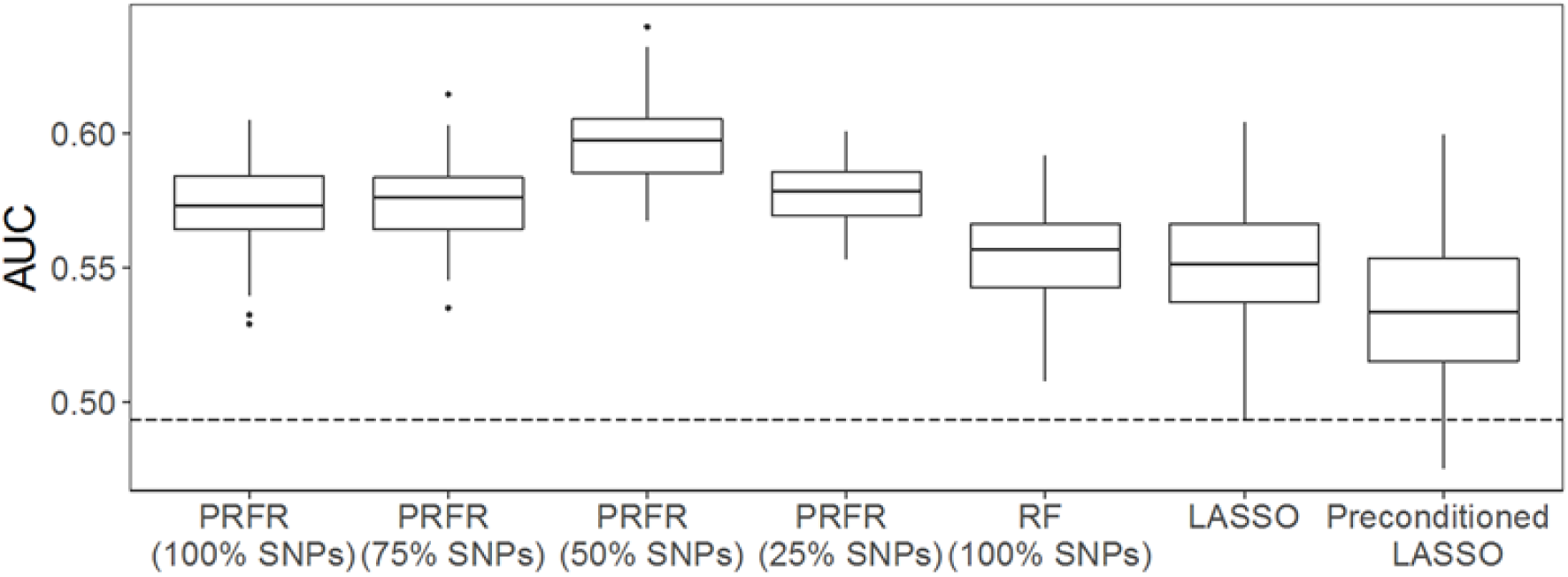
Comparison of prediction areas under the curve (AUCs) on the hold-out validation dataset between different multivariate radiation-associated contralateral breast cancer models. Each box plot indicates the fluctuation of AUCs over 100 iterations of 5-fold cross validation. *Abbreviations*: LASSO - least absolute shrinkage and selection operator; PRFR - preconditioned random forest regression; RF - random forest; SNPs - single nucleotide polymorphisms.

### Biological correlates and plausibility

In the analysis using subsets of the SNPs, the validation AUC of the PRFR model was the largest when only the top 50% of the SNPs were considered (Figure 3). This resulted in 356 SNPs, among which 149 SNPs were located in uninformative intergenic regions (out of the range of 50,000 base pairs from genes). The remaining 207 SNPs were mapped to 188 genes. The GSEA on the 188 genes resulted in 7 significant biological processes (Figure 4), forming 3 GO term groups, each containing more than one related GO term. The lowest *p*-value as a single term was observed for “cyclic adenosine monophosphate (cAMP)-mediated signaling” (GO:0019933, *p* = 2.6 × 10^-4^). This term, along with a term “second-messenger-mediated signaling” (GO:0019932, *p* = 2.8 × 10^-3^), formed a GO term group with the lowest group *p*-value of 2.6 × 10^-4^.

**Figure 4:**
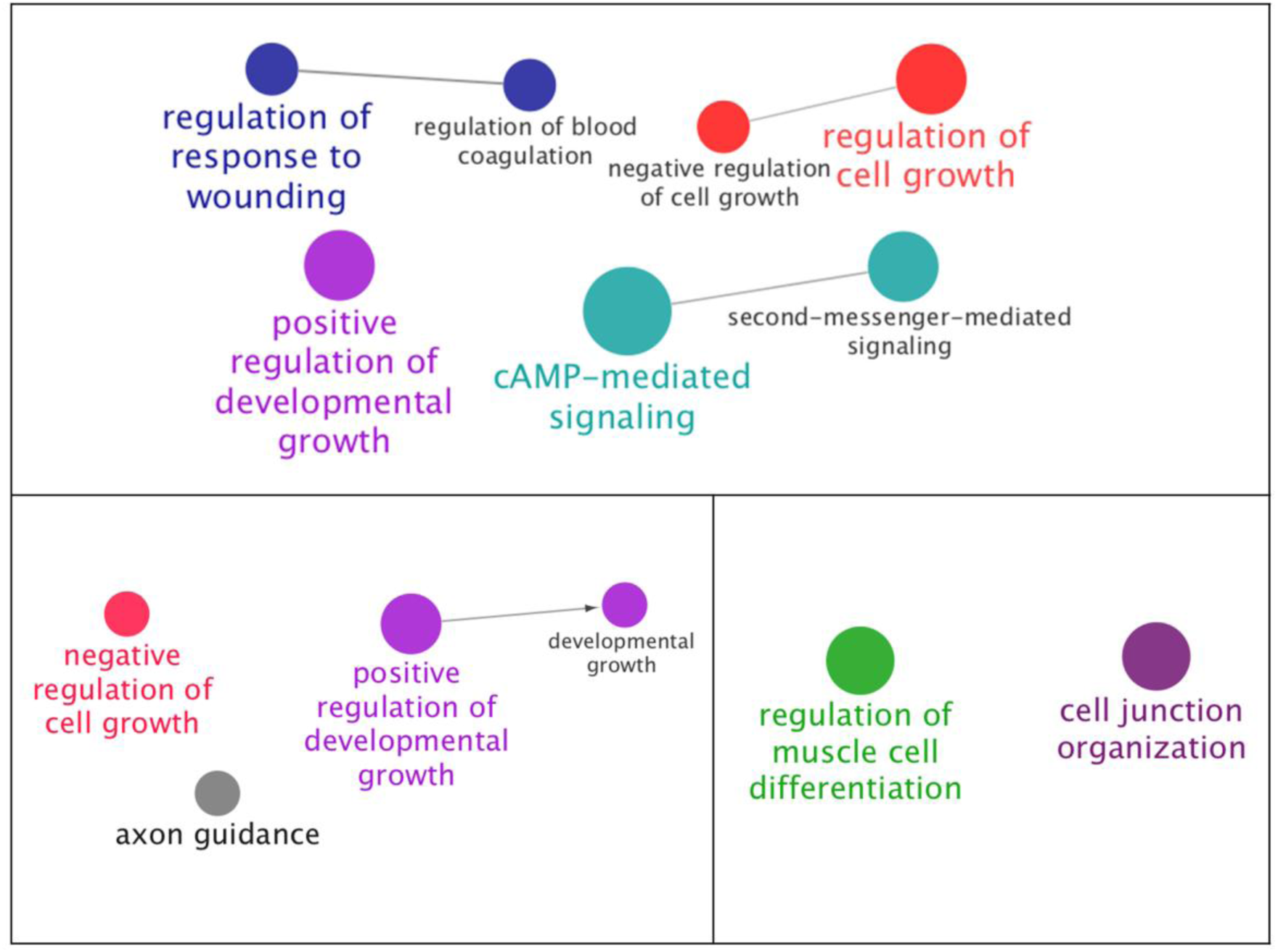
The biological processes that are significantly enriched from the 188 genes that were prioritized by preconditioned random forest regression (top), least absolute shrinkage and selection operator (bottom left), and univariate association strength (bottom right). The processes with lower p-values were drawn with larger node sizes.

Alternatively, when LASSO selection frequency was used to isolate the same number (188) of high-priority genes, 4 significant biological processes were observed. Two of them were in line with the results obtained from the PRFR-VIM prioritization (positive regulation of developmental growth and negative regulation of cell growth) (Figure 4). On the other hand, when a univariate *p*-value-based ranking was used, 2 GO terms were found to be significant, which did not overlap with those in the PRFR- or LASSO-based prioritization (Figure 4). Figure 5 shows the number of overlapping genes between different modeling methods as well as the overlaps between the SNPs that resulted in the 188 genes.

**Figure 5:**
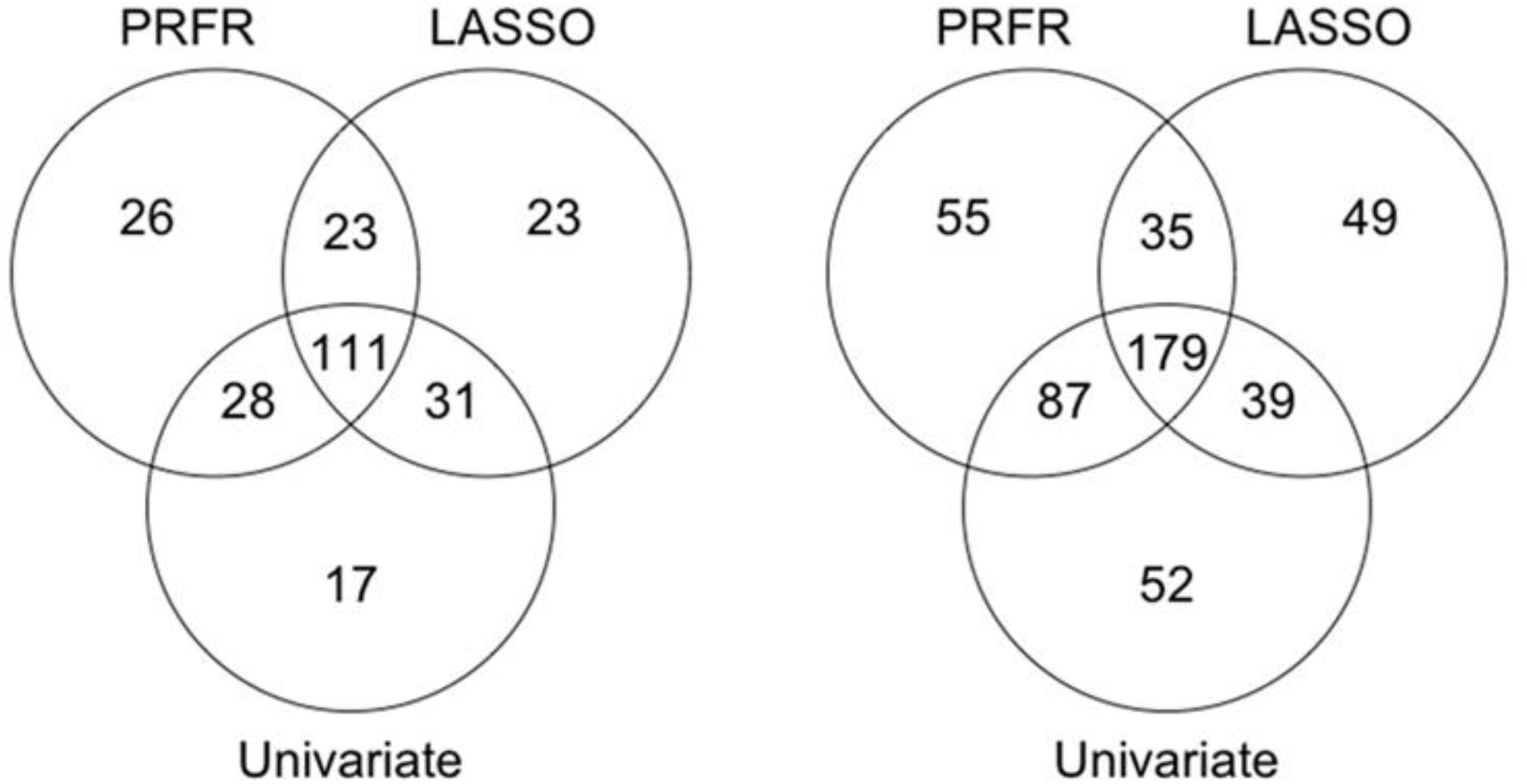
The number of overlapping genes between the 188 genes that were prioritized by 3 different methods (left; note that the univariate *p*-value ranking method consisted of 187 genes because many SNPs had the same *p*-values and inclusion of them exceeded 188 by a large amount) and the overlapping single nucleotide polymorphisms that resulted in the 188 genes (right). *Abbreviations*: PRFR - preconditioned random forest regression; LASSO - least absolute shrinkage and selection operator.

In an analysis of protein-protein interactions among the 188 genes from the PRFR model, MetaCore returned 8 interconnected proteins, forming 2 distinct clusters (Figure 6). The larger cluster contained 5 proteins, including B-cell lymphoma 6 (Bcl-6) that had connections with three proteins. The smaller cluster consisted of 3 proteins, where the Receptor tyrosine-protein kinase (ErbB)-4 was linked to Neuregulin 1 and 3. Separately, the literature survey indicated that all 8 proteins in the 2 clusters had previously been reported to be associated with breast cancer (Table 3). For 4 of the 8 proteins – CD63, Ephrin A, ERBB4, and Neuregulin 1 – the PubMed search found relevant publications linking the proteins to both radiation and carcinogenesis.

**Table 3.** Summary of literature survey on relevance of the genes/proteins in the protein-protein interaction network to breast cancer or radiation-associated carcinogenesis.

| Protein name | Summary of evidence |  |
| --- | --- | --- |
|  | Breast cancer carcinogenesis | Radiation-associated carcinogenesis |
| Bcl-6 | Highly expressed in breast cancer cells <sup>49</sup>; might promote invasion, migration, and growth by inducing epithelial-mesenchymal transition <sup>50</sup> |  |
| CD63 | Belongs to exosome family that plays a role in breast cancer progression <sup>51</sup> and malignancy <sup>52</sup> | Its secretion from head and neck carcinoma is increased after irradiation <sup>53</sup> |
| PTPRO | Inhibits ERBB2-driven breast carcinogenesis <sup>54</sup>; implicated in tamoxifen sensitivity <sup>55</sup> |  |
| BACH2 | DNA methylation marker for pathogenicity of a breast cancer subtype <sup>56</sup> |  |
| Ephrin A | EphA/Ephrin signaling pathway promotes breast cancer growth <sup>57</sup> or angiogenesis <sup>58</sup> | Ephrin signaling pathway responds to irradiation in an ATM-dependent manner <sup>46</sup> |
| ERBB4 | Activates growth-related downstream pathways, including MAPK kinase <sup>59</sup>; overexpression is associated with better prognosis <sup>60</sup> | Upon irradiation, triggers downstream proliferative pathways such as MAPK <sup>47</sup> |
| Neuregulin 1 | Promotes tumor growth via ERBB activation <sup>61</sup>, promoting cell migration <sup>62</sup> | Binds to and activates ERBB4, generating radioprotective signals <sup>48</sup> |
| Neuregulin 3 | Binds to and activates ERBB4 <sup>63</sup> |  |

**Figure 6:**
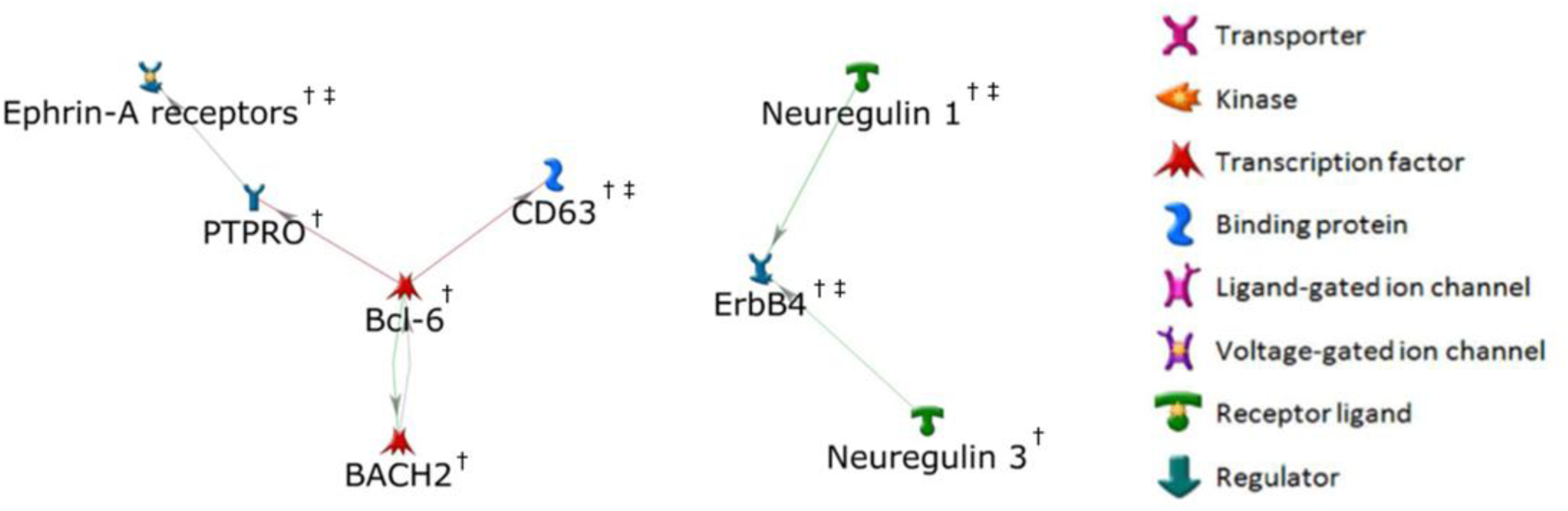
Two connected components among the 188 genes of high importance. †: associated with breast cancer, ‡: associated with radiation-induced carcinogenesis.

## Discussion

Traditional GWAS methods have focused on finding SNPs with main effects large enough to reach genome-wide significance (typically *p* < 5 × 10^-8^). However, any such genome-wide significant SNPs are typically not enough to account for the range of biological mechanisms responsible for the phenotype. The use of non-linear machine learning methods may allow for a more natural ranking of genetic importance within a many-SNP interaction paradigm. Our methodology serves dual purposes: 1) the non-linear multi-SNP models appear to yield better predictions of risk and thus can aid in clinical decision-making, and 2) these methods can be used to deduce possible biological mechanisms for the phenotype from the commonalities in known biology among the SNPs. Nonetheless, a fair number of SNPs used in the models are expected to be false positives under the relaxed *p*-value cutoff of 0.001. However, as seen in our results, the process of ranking SNPs using variable importance within the RF modeling could help to select SNPs for identifying more biologically relevant biomarkers, as has been suggested in previous studies ^41, 37^.

This work focused on a subgroup of the WECARE I Study that consisted of young women (≤ 40 years of age) who received a moderate level of scatter radiation dose to the contralateral breast (> 1 Gy). Using agnostic machine learning algorithms and bioinformatics tools that map to prior biological knowledge, a group of biological correlates were discovered, which are linked to breast carcinogenesis, with some links to pathways affecting radiation response.

The four biological process groups identified (Figure 4) were partly aligned with the biological mechanisms known to be pertinent to radiation response, including cAMP-mediated signaling and regulation of response to wounding (GO:1903034, *p* = 7.8 × 10^-3^). The most significant GO term in the analysis, cAMP-mediated signaling, has been shown to promote radiation-induced apoptosis in human lung cancer cells via interaction with the ATM gene ^42^. The regulation of response to wounding has been previously identified by the transcriptome experiment as a significantly enriched process in bystander cells adjacent to the region irradiated with alpha particles ^43^.

In addition, the genes for important SNPs were mapped onto the known network of human protein-protein interactions to identify key connected components, based on the premise that proteins that interact with each other tend to possess similar biological functions (Figure 6). The literature survey indicated that all the proteins in the two connected components were known to be involved in breast cancer. Moreover, in the GSEA results, significant biological processes pertaining to cell growth were revealed (Figure 4). These results corroborate the study by ^27^, indicating the overlap of genetic risk factors for RCBC with those for breast cancer risk among the general population. This is consistent with a mechanistic viewpoint that genetic defects that increase cancer risk might also be involved in radiosensitivity (e.g., DNA repair, cell cycle checkpoint control) ^44^. Interestingly, some of the clustered proteins have been identified as radiation responders or modifiers of radiation response. For example, the transcription regulator protein BACH2 has been shown to be down-regulated after ultraviolet radiation ^45^. More directly, Ephrin A, ERBB4, and Neuregulins have been identified for their potential roles as radiation-associated carcinogenesis biomarkers. Cheema et al. ^46^ showed that the Ephrin signaling pathway responds to irradiation in an ATM-dependent manner. ERBB4 is part of the epidermal growth factor receptor pathway that is activated by radiation, resulting in downstream effects on MAPK and cell proliferation ^47^. Neuregulin-1 binds to and activates ERBB4, providing radio-protective signals ^48^.

The impact of selecting between PRFR and LASSO methods on biological interpretation was assessed. LASSO is designed to find a parsimonious linear weighted sum of predictor variables. On the other hand, RF makes no prior assumption in model structures and is more flexible in accounting for interaction effects between predictors. Due to explicit regularization, LASSO induces sparsity in the modeling and thus effectively uses a fewer SNPs than PRFR. However, the sparsity by LASSO did not turn out to be beneficial for prediction of RCBC compared with PRFR (Figure 3) because many potentially important SNPs and interactions may have dropped out. Despite this limitation, the majority of GO terms from the LASSO-based prioritization were related or overlapped with those in the PRFR model. However, the biological processes that are potentially radiation-related, such as cAMP signaling or response to wounding, were only found in the PRFR-based GSEA. It was also observed that gene prioritization by PRFR or LASSO resulted in greater enrichment in relevant processes compared with the univariate *p*-value ranking method. This might indicate that the SNPs with moderate main effect strength (*p* ~10^-3^) also play a role in this phenotype via interaction, which would have been masked by applying a more stringent univariate cutoff. Taken together, the results support the concept that RF machine learning can be used not only as a prediction tool but also for identifying more interpretive biomarkers.

Planned future investigations, with additional genotype data from the second phase of the WECARE Study, will include comparison of important SNPs or gene sets between subgroups within the WECARE Study with different characteristics (age, radiation dose) as well as interaction effects between radiation dose and SNPs across the full range of dose to the contralateral breast.

### Conclusion

Using GWAS analysis combined with a non-linear machine learning approach, we created a prediction model of the risk of RCBC for young women who received a moderate level of radiation dose (> 1 Gy) to the contralateral breast. Although limited by sample size, the PRFR model achieved significant predictive performance in the hold-out validation and revealed plausible biological correlates associated with breast cancer or radiation response. This work demonstrates the potential of using machine learning and bioinformatics techniques for revealing the possible biological mechanisms of RCBC and for predicting risks to aid in clinical decision-making.

## Acknowledgments

This research was funded in part through National Institutes of Health/National Cancer Institute Cancer Center Support grant P30 CA008748, U01 CA083178, R01 CA097397, R01 CA114236, P01 CA196569, and the Breast Cancer Research Foundation grant BCRF-17-193.

We would like to thank Memorial Sloan Kettering Cancer Center (Coordinating Center) Investigators and Staff: Jonine L. Bernstein Ph.D. (WECARE Study P.I.); Marinela Capanu Ph.D.; Xiaolin Liang M.D.; Irene Orlow Ph.D.; Anne S. Reiner M.P.H.; Mark Robson M.D.; Meghan Woods M.P.H.

Collaborative Site Investigators: Leslie Bernstein Ph.D.; John D. Boice Jr. Sc.D.; Jennifer Brooks Ph.D.; Patrick Concannon Ph.D.; Dave V. Conti Ph.D.; David Duggan Ph.D.; Joanne W. Elena Ph.D., M.P.H.; Robert W. Haile Dr.P.H.; Esther M. John Ph.D.; Julia A. Knight Ph.D.; Charles F. Lynch M.D., Ph.D.; Kathleen E. Malone Ph.D.; Lene Mellemkjær Ph.D.; Jørgen H. Olsen M.D. DMSc.; Daniela Seminara Ph.D. M.P.H.; Roy E. Shore Ph.D., Dr.P.H.; Marilyn Stovall Ph.D.; Daniel O. Stram Ph.D.; Marc Tischkowitz M.D., Ph.D.; Duncan C. Thomas Ph.D. Collaborative Site Staff: Kristina Blackmore M.Sc.; Anh T. Diep; Judy Goldstein; Irene Harris B.S., C.M.D.; Rikke Langballe M.P.H.; Cecilia O'Brien; Susan Smith M.P.H.; Rita Weathers M.S.; Michele West Ph.D.

## References

1. Boice, J. D., Jr. Radiation and breast carcinogenesis. Medical and pediatric oncology 36, 508–513, doi:10.1002/mpo.1122 (2001).

2. Mertens, W. C., Hilbert, V. & Makari-Judson, G. Contralateral breast cancer: factors associated with stage and size at presentation. The breast journal 10, 304–312, doi:10.1111/j.1075-122X.2004.21333.x (2004).

3. Boice, J. D., Harvey, E. B., Blettner, M., Stovall, M. & Flannery, J. T. Cancer in the contralateral breast after radiotherapy for breast cancer. N Engl J Med 326, 781–785, doi:10.1056/NEJM199203193261201 (1992).

4. Preston, D. L. et al. Radiation effects on breast cancer risk: a pooled analysis of eight cohorts. Radiat Res 158, 220–235 (2002).

5. Largent, J. A. et al. Reproductive history and risk of second primary breast cancer: the WECARE study. Cancer Epidemiol Biomarkers Prev 16, 906–911, doi:10.1158/1055-9965.EPI-06-1003 (2007).

6. Brooks, J. D. et al. Reproductive status at first diagnosis influences risk of radiation-induced second primary contralateral breast cancer in the WECARE study. Int J Radiat Oncol Biol Phys 84, 917–924, doi:10.1016/j.ijrobp.2012.01.047 (2012).

7. Sisti, J. S. et al. Reproductive factors, tumor estrogen receptor status and contralateral breast cancer risk: results from the WECARE study. SpringerPlus 4, 825, doi:10.1186/s40064-015-1642-y (2015).

8. Ricceri, F. et al. Risk of second primary malignancies in women with breast cancer: Results from the European prospective investigation into cancer and nutrition (EPIC). Int J Cancer 137, 940–948, doi:10.1002/ijc.29462 (2015).

9. Chen, Y., Thompson, W., Semenciw, R. & Mao, Y. Epidemiology of contralateral breast cancer. Cancer epidemiology, biomarkers & prevention : a publication of the American Association for Cancer Research, cosponsored by the American Society of Preventive Oncology 8, 855–861 (1999).

10. Langballe, R. et al. Systemic therapy for breast cancer and risk of subsequent contralateral breast cancer in the WECARE Study. Breast Cancer Res 18, 65, doi:10.1186/s13058-016-0726-0 (2016).

11. Davies, C. et al. Relevance of breast cancer hormone receptors and other factors to the efficacy of adjuvant tamoxifen: patient-level meta-analysis of randomised trials. Lancet 378, 771–784, doi:10.1016/s0140-6736(11)60993-8 (2011).

12. Hill-Kayser, C. E., Harris, E. E., Hwang, W. T. & Solin, L. J. Twenty-year incidence and patterns of contralateral breast cancer after breast conservation treatment with radiation. Int J Radiat Oncol Biol Phys 66, 1313–1319, doi:10.1016/j.ijrobp.2006.07.009 (2006).

13. Hemminki, K., Ji, J. & Forsti, A. Risks for familial and contralateral breast cancer interact multiplicatively and cause a high risk. Cancer Res 67, 868–870, doi:10.1158/0008-5472.can-06-3854 (2007).

14. Reiner, A. S. et al. Breast Cancer Family History and Contralateral Breast Cancer Risk in Young Women: An Update From the Women’s Environmental Cancer and Radiation Epidemiology Study. J Clin Oncol, JCO2017773424, doi:10.1200/JCO.2017.77.3424 (2018).

15. Stovall, M. et al. Dose to the contralateral breast from radiotherapy and risk of second primary breast cancer in the WECARE study. Int J Radiat Oncol Biol Phys 72, 1021–1030, doi:10.1016/j.ijrobp.2008.02.040 (2008).

16. Begg, C. B., Haile, R. W., Borg, Å. & et al. Variation of breast cancer risk among brca1/2 carriers. Jama 299, 194–201, doi:10.1001/jama.2007.55-a (2008).

17. Graeser, M. K. et al. Contralateral breast cancer risk in BRCA1 and BRCA2 mutation carriers. J Clin Oncol 27, 5887–5892, doi:10.1200/jco.2008.19.9430 (2009).

18. Bernstein, J. L. et al. Radiation exposure, the ATM Gene, and contralateral breast cancer in the women’s environmental cancer and radiation epidemiology study. Journal of the National Cancer Institute 102, 475–483, doi:10.1093/jnci/djq055 (2010).

19. Brooks, J. D. et al. Variants in activators and downstream targets of ATM, radiation exposure, and contralateral breast cancer risk in the WECARE study. Human mutation 33, 158–164, doi:10.1002/humu.21604 (2012).

20. Bernstein, J. L. et al. Contralateral breast cancer after radiotherapy among BRCA1 and BRCA2 mutation carriers: a WECARE study report. Eur J Cancer 49, 2979–2985, doi:10.1016/j.ejca.2013.04.028 (2013).

21. Renwick, A. et al. ATM mutations that cause ataxia-telangiectasia are breast cancer susceptibility alleles. Nat Genet 38, 873–875, doi:10.1038/ng1837 (2006).

22. Antoniou, A. C. et al. A locus on 19p13 modifies risk of breast cancer in BRCA1 mutation carriers and is associated with hormone receptor-negative breast cancer in the general population. Nat Genet 42, 885–892, doi:10.1038/ng.669 (2010).

23. Cybulski, C. et al. Risk of breast cancer in women with a CHEK2 mutation with and without a family history of breast cancer. J Clin Oncol 29, 3747–3752, doi:10.1200/jco.2010.34.0778 (2011).

24. Michailidou, K. et al. Association analysis identifies 65 new breast cancer risk loci. Nature 551, 92–94, doi:10.1038/nature24284 (2017).

25. Takahashi, M. et al. The FOXE1 locus is a major genetic determinant for radiation-related thyroid carcinoma in Chernobyl. Hum. Mol. Genet. 19, 2516–2523, doi:10.1093/hmg/ddq123 (2010).

26. Best, T. et al. Variants at 6q21 implicate PRDM1 in the etiology of therapy-induced second malignancies after Hodgkin’s lymphoma. Nat Med 17, 941–943, doi:10.1038/nm.2407 (2011).

27. Robson, M. E. et al. Association of Common Genetic Variants With Contralateral Breast Cancer Risk in the WECARE Study. JNCI: Journal of the National Cancer Institute 109, djx051–djx051, doi:10.1093/jnci/djx051 (2017).

28. Bernstein, J. L. et al. Study design: evaluating gene-environment interactions in the etiology of breast cancer - the WECARE study. Breast cancer research : BCR 6, R199–214, doi:10.1186/bcr771 (2004).

29. Oh, J. H. et al. Computational methods using genome-wide association studies to predict radiotherapy complications and to identify correlative molecular processes. Scientific reports 7, 43381, doi:10.1038/srep43381 (2017).

30. Lee, S. et al. Machine Learning on a Genome-wide Association Study to Predict Late Genitourinary Toxicity After Prostate Radiation Therapy. Int J Radiat Oncol Biol Phys, doi:10.1016/j.ijrobp.2018.01.054 (2018).

31. Howie, B., Marchini, J. & Stephens, M. Genotype imputation with thousands of genomes. G3 1, 457–470, doi:10.1534/g3.111.001198 (2011).

32. Howie, B., Fuchsberger, C., Stephens, M., Marchini, J. & Abecasis, G. R. Fast and accurate genotype imputation in genome-wide association studies through pre-phasing. Nat. Genet. 44, 955–959, doi:10.1038/ng.2354 (2012).

33. Reiner, A. S. et al. Risk of asynchronous contralateral breast cancer in noncarriers of BRCA1 and BRCA2 mutations with a family history of breast cancer: a report from the Women’s Environmental Cancer and Radiation Epidemiology Study. J Clin Oncol 31, 433–439, doi:10.1200/jco.2012.43.2013 (2013).

34. Bertelsen, L. et al. Effect of systemic adjuvant treatment on risk for contralateral breast cancer in the Women’s Environment, Cancer and Radiation Epidemiology Study. Journal of the National Cancer Institute 100, 32–40, doi:10.1093/jnci/djm267 (2008).

35. Hoh, J. et al. Selecting SNPs in two-stage analysis of disease association data: a model-free approach. Ann Hum Genet 64, 413–417 (2000).

36. Chen, X. & Ishwaran, H. Random forests for genomic data analysis. Genomics 99, 323–329, doi:10.1016/j.ygeno.2012.04.003 (2012).

37. Nguyen, T. T., Huang, J., Wu, Q., Nguyen, T. & Li, M. Genome-wide association data classification and SNPs selection using two-stage quality-based Random Forests. BMC genomics 16 Suppl 2, S5, doi:10.1186/1471-2164-16-S2-S5 (2015).

38. The Gene Ontology Consortium. Expansion of the Gene Ontology knowledgebase and resources. Nucleic Acids Res 45, D331–d338, doi:10.1093/nar/gkw1108 (2017).

39. Bindea, G. et al. ClueGO: a Cytoscape plug-in to decipher functionally grouped gene ontology and pathway annotation networks. Bioinformatics 25, 1091–1093, doi:10.1093/bioinformatics/btp101 (2009).

40. Zheng, G., Freidlin, B. & Gastwirth, J. L. Robust genomic control for association studies. Am J Hum Genet 78, 350–356, doi:10.1086/500054 (2006).

41. Lunetta, K. L., Hayward, L. B., Segal, J. & Van Eerdewegh, P. Screening large-scale association study data: exploiting interactions using random forests. BMC genetics 5, 32, doi:10.1186/1471-2156-5-32 (2004).

42. Cho, E. A., Kim, E. J., Kwak, S. J. & Juhnn, Y. S. cAMP signaling inhibits radiation-induced ATM phosphorylation leading to the augmentation of apoptosis in human lung cancer cells. Molecular cancer 13, 36, doi:10.1186/1476-4598-13-36 (2014).

43. Yeles, C. et al. Integrative Bioinformatic Analysis of Transcriptomic Data Identifies Conserved Molecular Pathways Underlying Ionizing Radiation-Induced Bystander Effects (RIBE). Cancers (Basel) 9, doi:10.3390/cancers9120160 (2017).

44. International Commission on Radiological Protection. Genetic susceptibility to cancer: ICRP publication 79. Ann ICRP 28, 1–157 (1998).

45. Uittenboogaard, L. M. et al. BACH2: a marker of DNA damage and ageing. DNA repair 12, 982–992, doi:10.1016/j.dnarep.2013.08.016 (2013).

46. Cheema, A. K. et al. Functional proteomics analysis to study ATM dependent signaling in response to ionizing radiation. Radiat Res 179, 674–683, doi:10.1667/RR3198.1 (2013).

47. Bowers, G. et al. The relative role of ErbB1-4 receptor tyrosine kinases in radiation signal transduction responses of human carcinoma cells. Oncogene 20, 1388–1397, doi:10.1038/sj.onc.1204255 (2001).

48. Dent, P. et al. Stress and radiation-induced activation of multiple intracellular signaling pathways. Radiat Res 159, 283–300 (2003).

49. Walker, S. R. et al. The transcriptional modulator BCL6 as a molecular target for breast cancer therapy. Oncogene 34, 1073–1082, doi:10.1038/onc.2014.61 (2015).

50. Yu, J. M. et al. BCL6 induces EMT by promoting the ZEB1-mediated transcription repression of E-cadherin in breast cancer cells. Cancer Lett 365, 190–200, doi:10.1016/j.canlet.2015.05.029 (2015).

51. Shi, J., Ren, Y., Zhen, L. & Qiu, X. Exosomes from breast cancer cells stimulate proliferation and inhibit apoptosis of CD133+ cancer cells in vitro. Molecular medicine reports 11, 405–409, doi:10.3892/mmr.2014.2749 (2015).

52. Tominaga, N. et al. RPN2-mediated glycosylation of tetraspanin CD63 regulates breast cancer cell malignancy. Molecular cancer 13, 134, doi:10.1186/1476-4598-13-134 (2014).

53. Jelonek, K. et al. Ionizing radiation affects protein composition of exosomes secreted in vitro from head and neck squamous cell carcinoma. Acta biochimica Polonica 62, 265–272, doi:10.18388/abp.2015_970 (2015).

54. Dong, H. et al. PTPRO represses ERBB2-driven breast oncogenesis by dephosphorylation and endosomal internalization of ERBB2. Oncogene 36, 410–422, doi:10.1038/onc.2016.213 (2017).

55. Ramaswamy, B. et al. Estrogen-mediated suppression of the gene encoding protein tyrosine phosphatase PTPRO in human breast cancer: mechanism and role in tamoxifen sensitivity. Molecular endocrinology 23, 176–187, doi:10.1210/me.2008-0211 (2009).

56. Flower, K. J. et al. DNA methylation profiling to assess pathogenicity of BRCA1 unclassified variants in breast cancer. Epigenetics 10, 1121–1132, doi:10.1080/15592294.2015.1111504 (2015).

57. Kaenel, P., Mosimann, M. & Andres, A. C. The multifaceted roles of Eph/ephrin signaling in breast cancer. Cell adhesion & migration 6, 138–147, doi:10.4161/cam.20154 (2012).

58. Brantley-Sieders, D. M. et al. Impaired tumor microenvironment in EphA2-deficient mice inhibits tumor angiogenesis and metastatic progression. FASEB journal : official publication of the Federation of American Societies for Experimental Biology 19, 1884–1886, doi:10.1096/fj.05-4038fje (2005).

59. Sundvall, M. et al. Role of ErbB4 in breast cancer. Journal of mammary gland biology and neoplasia 13, 259–268, doi:10.1007/s10911-008-9079-3 (2008).

60. Thor, A. D., Edgerton, S. M. & Jones, F. E. Subcellular localization of the HER4 intracellular domain, 4ICD, identifies distinct prognostic outcomes for breast cancer patients. The American journal of pathology 175, 1802–1809, doi:10.2353/ajpath.2009.090204 (2009).

61. Tsai, M. S., Shamon-Taylor, L. A., Mehmi, I., Tang, C. K. & Lupu, R. Blockage of heregulin expression inhibits tumorigenicity and metastasis of breast cancer. Oncogene 22, 761–768, doi:10.1038/sj.onc.1206130 (2003).

62. Haskins, J. W., Nguyen, D. X. & Stern, D. F. Neuregulin 1-activated ERBB4 interacts with YAP to induce Hippo pathway target genes and promote cell migration. Science signaling 7, ra116, doi:10.1126/scisignal.2005770 (2014).

63. Kogata, N., Zvelebil, M. & Howard, B. A. Neuregulin 3 and erbb signalling networks in embryonic mammary gland development. Journal of mammary gland biology and neoplasia 18, 149–154, doi:10.1007/s10911-013-9286-4 (2013).

